# Ecological specialization and local adaptation in sympatric sexual and asexual grass thrips species

**DOI:** 10.1101/2023.12.01.569600

**Authors:** Karim Ghali, Casper J. van der Kooi, Elodie Ramella, Morgane Henry, Tanja Schwander

## Abstract

The maintenance of sex is difficult to explain in the face of the demographic advantages of asexuality, especially when sexual and asexual lineages co-occur and compete. Here, we test if niche divergence and specialization can contribute to the maintenance of sympatric populations of two closely related, sexual and asexual *Aptinothrips* grass thrips species. In mesocosm experiments, ecological niches and ecological specialization were inferred from thrips performances on different grass species used as hosts in natural populations. Sexual and asexual thrips performed best on different grass hosts, indicating niche differentiation. The asexual species was also characterized by a broader fundamental ecological niche than the sexual one. However, niche differentiation is unlikely to explain the maintenance of the two species in sympatry because the reproductive rate of asexual females generally exceeded that of sexual ones. Surprisingly, the asexual but not sexual species showed geographic variation in the fundamental niche. This geographic variation likely stems from different clonal assemblages at different locations because different asexual genotypes have different ecological niches. Across natural populations, the performance of asexual females on a specific grass species was further positively correlated with the frequency of that grass species, consistent with adaptation to locally abundant grasses. Altogether, our results suggest that niche differentiation contributes little to the maintenance of grass thrips species with different reproductive modes and that asexuality facilitates adaptation to a diversity of co-occurring host plants.

## Introduction

The ability to reproduce is a key trait distinguishing living organisms from inorganic matter, yet reproductive strategies vary widely among organisms (Bell 1982; Suomalainen et al. 1987). Which factors contribute to the maintenance of alternative reproductive strategies in natural populations? This question has been particularly difficult to solve in the context of sexual versus asexual reproduction. Despite many studies focusing on the (dis)advantages of sexual and asexual reproduction, our knowledge of the factors maintaining both reproductive strategies in natural populations remains limited (Bell 1982; Case and Taper 1986; Neiman et al. 2018).

An example of co-occurring species and lineages with different reproductive modes is found in *Aptinothrips* grass thrips, which are small (∼1 mm), wingless insects found in grasslands throughout the Holarctic region of the world (Palmer 1975). The species *A. elegans* is sexual, *A. karnyi* and *A. stylifer* are asexual, and *A. rufus* comprises both sexual and asexual lineages (van der Kooi and Schwander 2014; Fontcuberta García-Cuenca et al. 2016). Species and lineages with different reproductive modes co-occur frequently in the same meadows in natural populations. Under laboratory conditions in mesocosms however, asexual lineages consistently outcompete their sexual relatives (Lavanchy et al. 2016).

We previously evaluated whether habitat heterogeneity contributes to the maintenance of species and lineages with different reproductive modes (Lavanchy et al. 2016), given the potential benefits of sex are predicted to increase with habitat heterogeneity (Lenormand 2002; Agrawal 2009). *Aptinothrips* depend on grasses (Poaceae) for their entire life cycle, as grasses provide resources, shelter and egg laying substrates (Palmer 1975). Thus, a key component for *Aptinothrips* habitat heterogeneity is the diversity of local grass species. Contrary to predictions, in natural grass meadows experimentally manipulated for increased and decreased grass diversity, we found that increased habitat diversity consistently favored asexual over sexual *Aptinothrips* species (Lavanchy et al. 2016). Because different asexual clones vary in their performance on different grass hosts (van der Kooi et al. 2019), we hypothesized that an increased diversity of host plants would accommodate a broader panel of clones, or, more generally, that the composition of local grass species communities would drive the composition of clone genotypes.

Here, we follow up on these previous studies by testing whether the maintenance of closely related, sympatric *Aptinothrips* lineages with different reproductive modes is facilitated by differences in niche strategy and niche size among lineages. We focus on the two lineages with the broadest distribution overlap: *A. elegans* (sexual) and asexual lineages of *A. rufus*. To characterize niche strategies and sizes, we measure the reproductive success of individual females on different grass species that are abundant in natural thrips habitats. We test whether the two species are characterized by different fundamental niches and/or different levels of ecological specialization and whether niche strategies vary geographically. Because we find geographic variation for niche strategies in the asexual but not sexual species, we further measure the performance of asexual females originating from different grass communities to test whether a higher abundance of a specific grass species translates into higher performances of the local thrips on that grass species.

## Methods

### Evaluating fundamental niche differences between grass thrips collected from different grass communities

We chose grass meadows from three locations in western Switzerland where we expected to find different grass communities based on different elevation and exposition: Lausanne (hereafter: UNIL, 405 m elevation), La Sarraz (559 m) and Blonay (953 m). From each of these locations, we then collected adult female grass thrips during September-November 2015 and, simultaneously, standardized samples of grass plants to genetically characterize the grass community composition (see Supplementary methods; Table 1). Thrips were collected by randomly beating grasses above a white tray, and adult *Aptinothrips* females, recognizable by their body movements and shape/size, were transferred with a paintbrush to a 50mL Falcon tube with freshly cut grass pieces until identification in the laboratory. This collection strategy mixes all thrips collected from different grass hosts, such that it is not possible to know which individual thrips used which specific grass host in the field. Note that in our previous studies (van der Kooi and Schwander 2014; Fontcuberta García-Cuenca et al. 2016; Lavanchy et al. 2016), adult females of sexual species and lineages collected in the field were always mated, given they always produced male and female offspring (virgin sexual females only produce males, given the haplo-diploid sex determination system in thrips). In the laboratory, female *Aptinothrips* were then identified to the species level by sorting individuals on a CO_2_ anesthetization plate under a 50x magnification dissecting scope, following Zur Strassen (2003).

**Table 1:** Assessment of the relative coverage of grass species at the three sampling locations. The grass species used in the laboratory mesocosms to assess thrips performance are indicated

| Grass species | Location |  |  |
| --- | --- | --- | --- |
|  | UNIL | La Sarraz | Blonay |
| *Agropyron repens* | 0 | 14 | 0 |
| *Agrostis capillaris* | 0 | 0.33 | 0 |
| *Anthoxanthum odoratum* | 5.67 | 2 | 1.33 |
| *Arrhenatherum elatius* | 14.67 | 2 | 30.67 |
| *Avenula pubescens* | 3 | 0.33 | 0 |
| *Brachypodium pinnatum* | 0 | 0 | 1.33 |
| *Bromus erectus* | 10.33 | 6 | 38.67 |
| *Carex caryophyllea* | 0 | 9.33 | 0 |
| *Carex flacca* | 0 | 0 | 0.33 |
| ***Dactylis glomerata*** | 19 | 0 | 4 |
| *Festuca arundinacea* | 0 | 0 | 5.33 |
| ***Festuca ovina*** | 3.33 | 27.33 | 4 |
| ***Festuca rubra*** | 0 | 0 | 6.33 |
| *Holcus lanatus* | 15.33 | 0 | 4 |
| ***Phleum pratense*** | 0.67 | 0 | 0 |
| ***Poa pratensis*** | 21 | 17 | 2.67 |
| *Poa trivialis* | 1 | 0 | 0 |
| *Trisetum flavescens* | 0 | 21.67 | 0 |

The performance of sexual *A. elegans* and asexual *A. rufus* females from the three locations was measured on seven grass species, with ten replicates per combination (2 species x 3 locations x 7 grass species x 10 replicates = 420 females in total). We chose seven grass species that are commonly found in natural habitats of *Aptinothrips* (Lavanchy et al. 2016) and for which seeds were commercially available: alpine bluegrass (*Poa alpina*), common meadow-grass (*Poa pratensis*), red fescue (*Festuca rubra*), sheep fescue (*Festuca ovina*), timothy-grass (*Phleum pratense*), orchard grass (*Dactylis glomerata*) and wheat (*Triticum aestivum*). However, we did not know the frequency of these species at the three surveyed locations *a priori*, as the composition of the grass species communities was determined after collecting the thrips. Thus, only five of the seven grass species used for the performance assays were present at the three thrips sampling locations. To measure thrips performances on each grass species, three-week old grass plantlets were placed in sealed 50 ml Falcon tubes for thrips breeding. Females were allowed to reproduce for 27 days (which corresponds to one generation) at 23°C during the day and 20°C during the night in a greenhouse with a relative humidity of 50% and an artificial light regime of L:D = 14/10 hours, added to the natural light conditions. After the 27 days, thrips were extracted from the plants via heat gradients on Berlese funnels, fixed in 70% ethanol, and adults and larvae from each replicate were counted under a dissecting scope at 50x magnification. Due to logistic constraints, the replicates of each treatment were divided into three batches, with a week separating the batches. Seven of the initially 420 replicates were lost because the grass plantlets died during the first week after transferring the thrips (for unknown reasons; four on *P. pratensis* from UNIL, two on *P. pratense* from La Sarraz and one on *P. pratense* from Blonay).

To test if the two *Aptinothrips* species were characterized by different niches, we used Linear Mixed Models (LMMs) as implemented in the R package *lme4* (Bates et al. 2015; R core team 2019). We tested if thrips species, grass species, sampling location, and all possible interactions affected the number of offspring produced. Batch was included as a random factor in the model. To test for significance, the number of offspring (response variable) was randomized within the three batches 100’000 times and F-values corresponding to main and interaction effects were extracted. The p-value of the factor of interest then corresponds to the proportion of F-values that were equal to or higher than the observed F-values.

### Niche breadth

We quantified the dietary niche breadth of *A. elegans* and *A. rufus* using the Shannon-Weaver index *H’* (Whittaker 1972), a measure of system entropy frequently used as a measure of dietary niche breadth (Sargeant 2007): 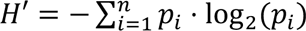. In this index, *n* is the number of grass species and *p_i_* the performances (i.e.: the number of offspring produced) on the *i*-th grass species. The index is then scaled to vary from 0 (extreme specialist) to 1 (extreme generalist). To compare the niche sizes between *A. elegans* and *A. rufus* overall and at each sampling location, we assessed whether the 95% confidence intervals (CIs) of the two groups overlapped. The number of offspring produced were randomly resampled 10’000x within species (with replacement) and random *H’* values were computed as above. Confidence intervals represent the 95% quantile of the random *H’* values.

These analyses revealed that the asexual species *A. rufus* was an extreme generalist across grass species within the Poaceae family (see results). Generalism in asexual populations can result from a combination of different clones with distinct, narrow niches, or directly from generalist clones with very broad niches (see discussion). To distinguish between these two mechanisms, we re-analyzed previously published performance data from nine genetically distinct clonal lines from (van der Kooi et al. 2019). These clonal lines were established from adult *A. rufus* females collected at the same three locations used in the present study, in February 2016. The performance data were generated via the same methodology as described above, for the six non-cultivated grass species (i.e., excluding wheat, *T. aestivum*) and with 7-14 replicates per clonal line and grass species (van der Kooi et al. 2019).

### Link between thrips performance and host grass species abundance

We found that the fundamental niches of asexual *A. rufus* varied among the three sampled locations UNIL, La Sarraz and Blonay (see results). Such variation could potentially be driven by selection for adaptation to locally abundant grass host species. To evaluate this possibility, we tested for a correlation between the abundance of two grass species (*F. rubra* and *P. trivialis*) and thrips performance across 16 grass meadows in the Swiss Prealps. The 16 meadows were selected to cover a broad range of abundances of the two grass species (*F. rubra*: ranging from 0% to 90%; *P. trivialis*: ranging from 0% to 38%), based on plant inventories conducted by (Dubuis et al. 2013). We collected and identified adult asexual *A. rufus* females from each of the 16 meadows as described above and measured the performance of at least 7 and up to 35 females from each meadow on each grass species (678 females in total, see Table S1 for details; note that *A. rufus* was very rare in four meadows and we therefore assessed the performance of *A. rufus* females on only one grass species for these meadows). We then tested for an association between grass abundance and thrips performance separately for the two grass species using an LMM with the date of collection (fifteen dates, Table S1) included as a random factor. To test for significance, thrips performance (i.e., the number of offspring produced; response variable) was randomized 100’000 times within the fifteen collection dates and F-values corresponding to main and interaction effects were extracted. The p-value of the factor of interest then corresponds to the proportion of F-values that were equal to or higher than the observed F-values.

## Results

Asexual *A. rufus* females produced significantly more offspring than sexual *A. elegans* females (permutation LMM; df=1, p<0.01, Figure 1; see also Table S2). The grass species (df=6, p<0.01) used for rearing and the sampling location (df=2, p=0.027) also had a significant influence on the number of offspring produced (Figure 1). In addition, significant interactions between sampling location and thrips species (df=2, p<0.01), grass species and thrips species (df=5, p=0.03), and a triple interaction between thrips species, grass species and location (df=10, p=0.016) were detected, which prompted us to analyze each location separately (Figure 1). In two out of three locations, niche strategies differed between sexual *A. elegans* and asexual *A. rufus* (as revealed by a significant thrips species by grass species interaction for UNIL (permutation LMM; df=6, p=0.012) and La Sarraz (df=6, p= 0.041). The same trend was not significant in Blonay (df=6, p=0.111; Table S2).

**Figure 1:**
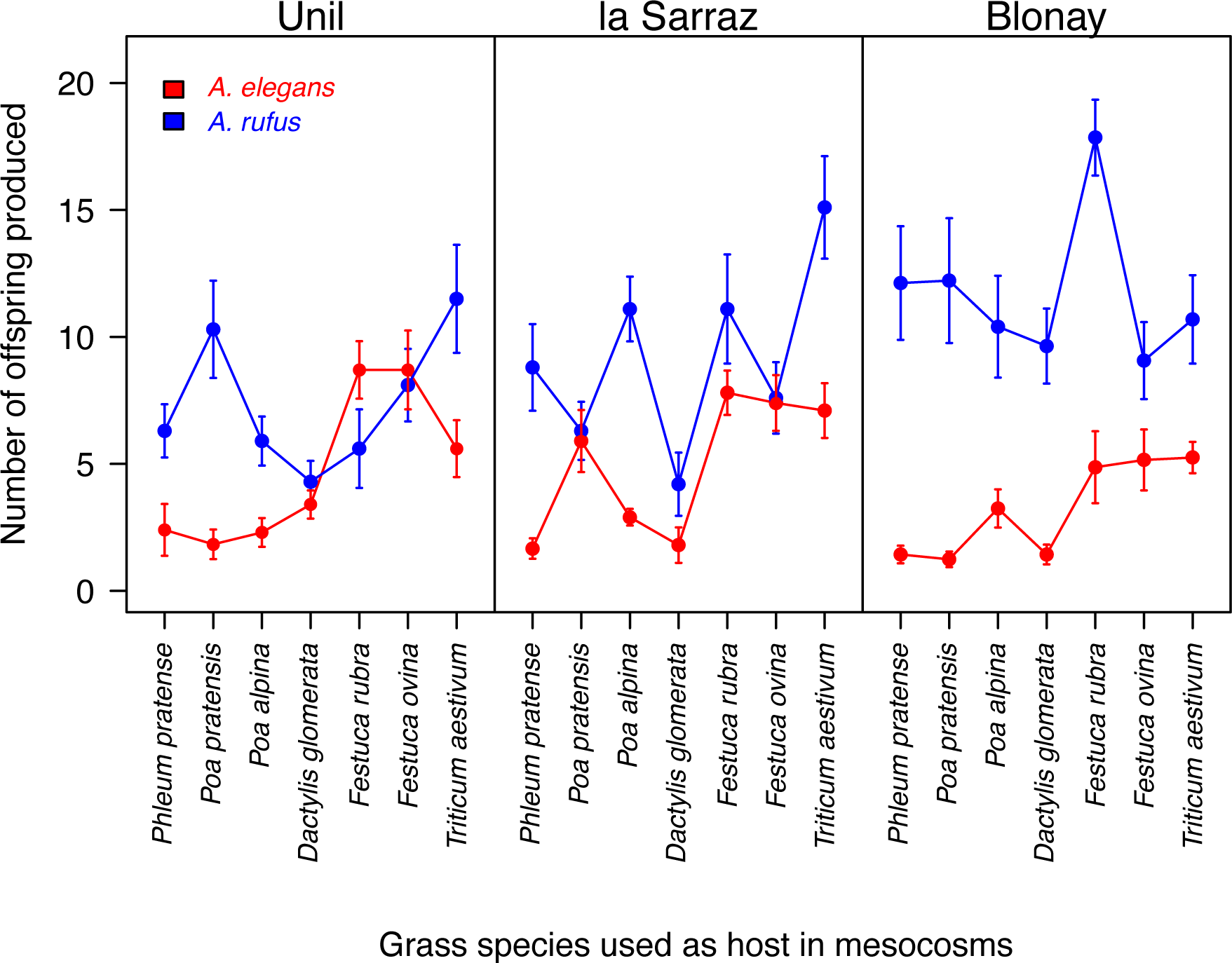
Average performance (number of offspring produced over 27 days) and standard deviation per grass species for sexual *A. elegans* and asexual *A. rufus* females for each sampled location. The lines connecting points have no biological meaning but are added for a better visualization.

*Aptinothrips* are specialists since they solely use grass species (Poaceae) as hosts, but within this plant family, both thrips species thrive on a variety of species i.e., they are generalists (Figure 2). Sexual *A. elegans* nevertheless had a somewhat narrower niche, that is, was significantly more specialized than the asexual species *A. rufus* across the three locations combined (*A. elegans*: *H’* = 0.93, 95% CI [0.89, 0.95], *A. rufus*: *H’* = 0.99, 95% CI [0.97, 0.99]; Figure 2). The species-level generalist phenotype for *A. rufus* was due to a combination of very generalist clones (clone 4 and 8, Figure 2) and more specialized ones (clone 2 and 9). The latter perform best on different grass species (*Festuca rubra* for clone 9 and *Poa pratensis* for clone 2; van der Kooi et al. 2019), meaning they are characterized by different fundamental niches.

**Figure 2:**
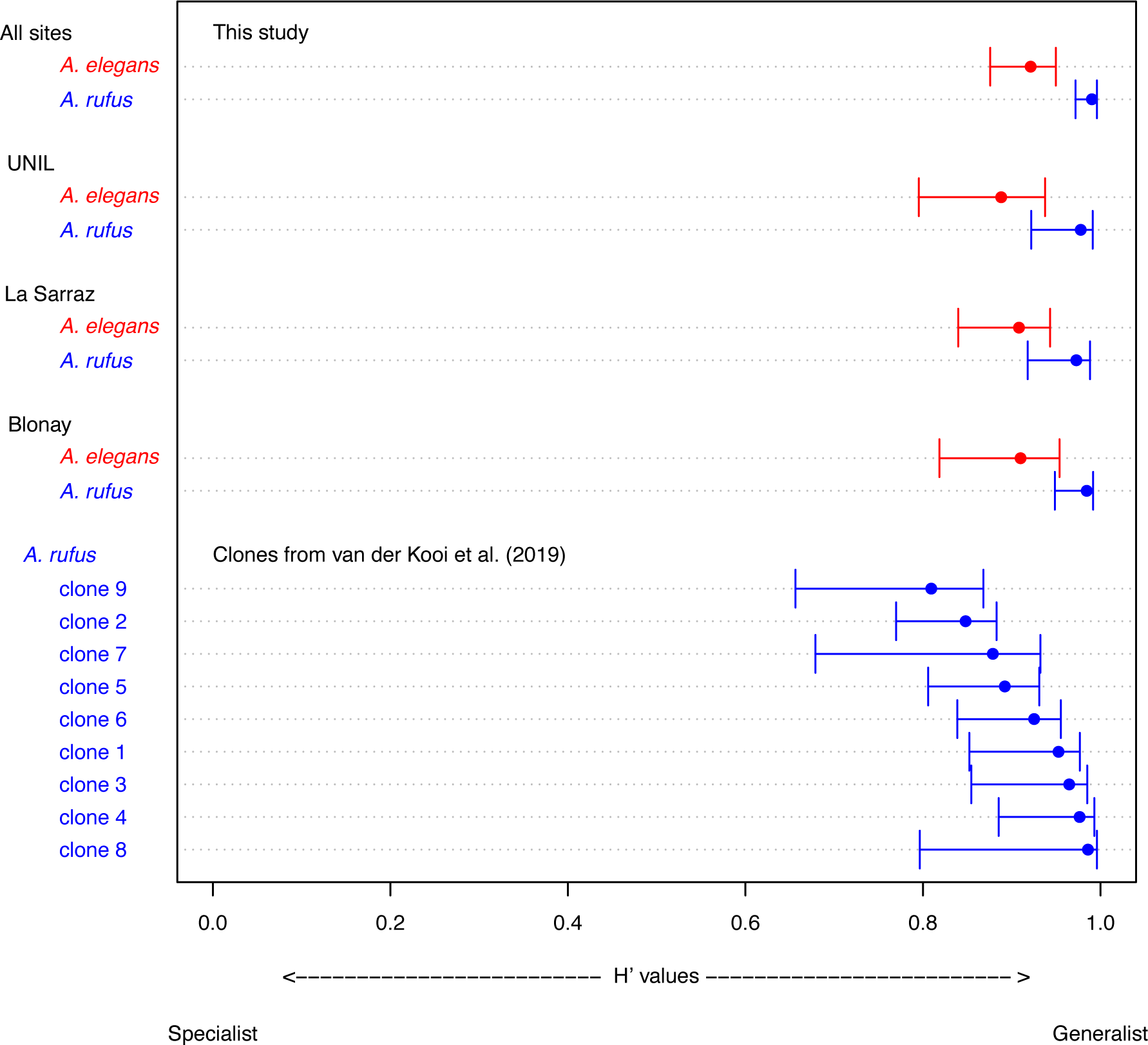
Shannon-Weaver indices (*H’*) for the sexual species *A. elegans* (red; *H’*: 0.93, CI 0.89-0.95) and the asexual species *A. rufus* (blue; *H’*: 0.99, CI 0.97-0.99) measured from performances on different grasses. The species-level index in the asexual species is further decomposed into clone-level indices. Confidence intervals are based on 10’000 bootstraps.

Sexual *A. elegans* from different locations exhibit similar fundamental niches. This is indicated by *A. elegans* females from different locations performing similarly across grass species (permutation LMM; df=12, p=0.26, Table S2, Figure S1). By contrast, the fundamental niche of asexual *A. rufus* varied geographically, as revealed by a significant interaction effect between grass species and location on thrips performance, permutation LMM; df=12, p=0.05, Table S2, Figure S1). The observed significant geographic variation in fundamental niche raises the question whether asexual *A. rufus* populations are locally adapted to abundant host grass species. If this is the case, a higher abundance of a certain grass species at a given location should translate into better performance of the local thrips on that grass species. For the grass *Festuca rubra* this was indeed the case: the more abundant *F. rubra* was at a specific location, the better the performance of asexual females on this species (permutation LMM; df=13, p<0.01, Figure 3; Table S3). However, there was no such relationship for *Poa trivialis* (df=13, p=0.943, Figure 3; Table S3), perhaps because this species was not present at high coverage at any of the 16 surveyed locations (maximum coverage 38%, Table S1).

**Figure 3:**
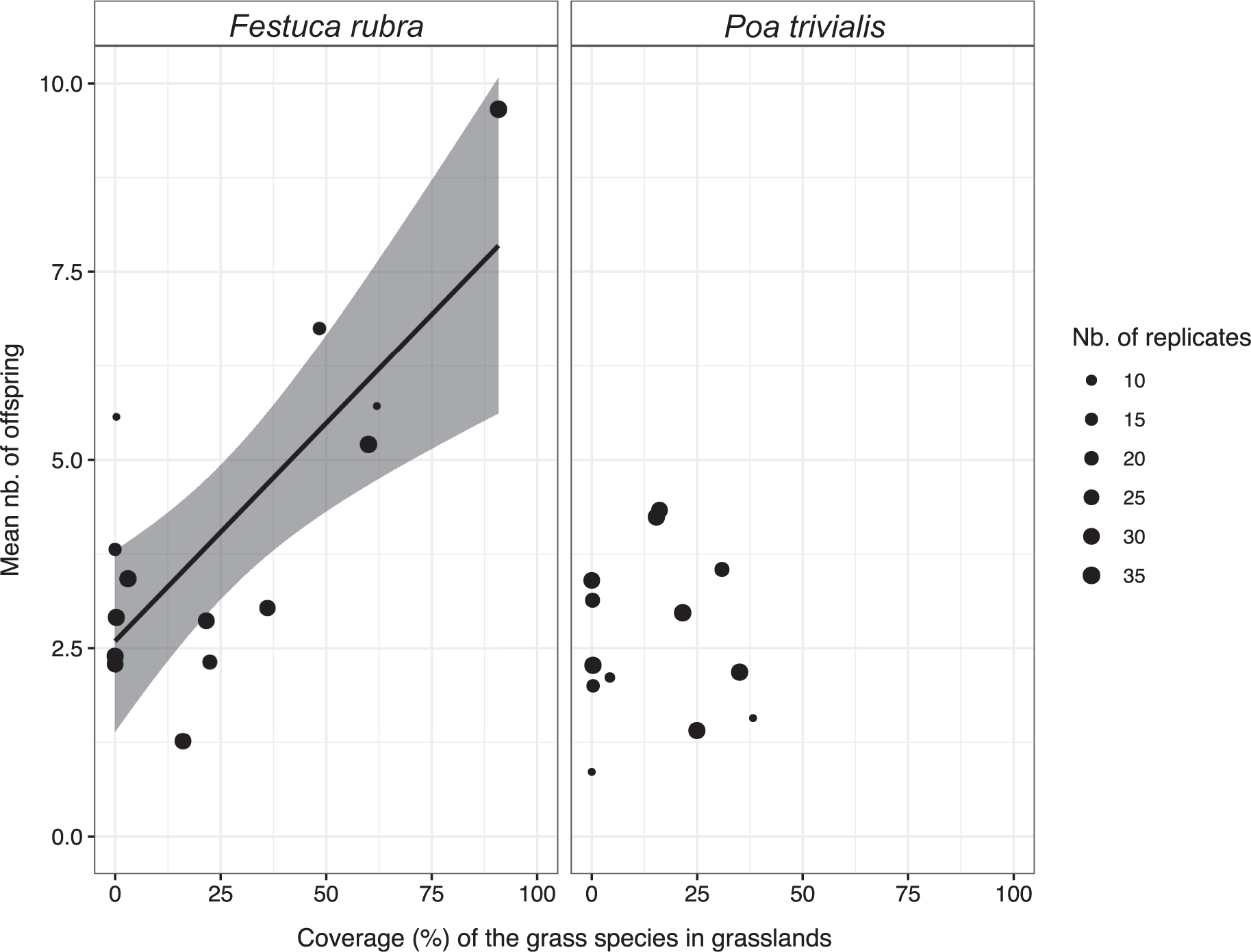
Correlation between the coverage of A) *Festuca rubra* or B) *Poa trivialis* in the field (%) and the performance of asexual thrips on those grass species. Each point represents one location, with thrips performance (number of offspring produced during 27 days) averaged across 7-35 females from that location. The line in panel A represents the LMM slope estimate, the gray shading shows the 95% confidence interval.

## Discussion

Sexual and asexual lineages often co-occur in the same geographic location, but whether they exhibit different niche strategies and whether potential niche differences may be sufficient to maintain sexual lineages in the face of competition from asexual ones remains largely unknown (but see O’Connell and Eckert 2001; Lehto and Haag 2010; Schmit et al. 2013; reviewed in Neiman et al. 2018). Here, we studied whether variation in ecological niches can contribute to the maintenance of *Aptinothrips* grass thrips species with different reproductive modes and which appear to compete in natural populations. We show that females of the asexual species *A. rufus* have a higher reproductive capacity than females of the sexual species *A. elegans*, corroborating previous findings in mesocosm habitats (Lavanchy et al. 2016). Asexual females produced 2.23 times more offspring than sexual females, which is added to the advantage of not producing males (Smith 1978; Avise 2008). Nevertheless, the two species systematically co-occur across grass meadows.

Despite the finding of different fundamental ecological niches between *A. rufus* and *A. elegans* at two of the three locations studied, the use of different host grass species could contribute to the co-existence of both species on *Festuca* grasses only at one location: the UNIL sampling location was the only population where sexual females produced more offspring than asexual ones. Sexual females from that location produced approximately 1.55 times more offspring than asexual ones if reared on *Festuca*. Given that the proportion of males among offspring in sexual *Aptinothrips* is typically below 0.3 (van der Kooi and Schwander 2014), this increase in offspring production is sufficient to compensate for the cost of male production in sexual species. Production of males is generally considered to be the most severe cost associated with sex (Lehtonen et al. 2012), although other costs, such as those linked to finding mates or mating, are typically difficult to quantify under natural conditions. Independent of the actual cost of sex in *Aptinothrips*, increased reproductive output of the sexual species on certain host plants could at least partially compensate for such costs. For example, *A. elegans* is associated with the presence of the grass species *Bromus erectus* in some natural populations of Switzerland (Lavanchy et al. 2016). Performances of *A. elegans* females on specific grass species also appear to be similar among locations (Figure S1), despite extensive variation in the frequency of these grasses at the sampling locations (Table 1). Adaptation of *A. elegans* to a reduced set of specific grass species might thus help explain why females of this species seem to be globally more productive on *Festuca* host species.

Contrary to *A. elegans*, females of asexual *A. rufus* showed variation in performances on different grass species according to their origin, raising the question of local adaptation. However, this pattern itself is not sufficient to conclude for an adaptation of *A. rufus* to local grass communities. Indeed, it may also be caused by specific life history traits. Individuals may be able to select their destination when dispersing and choose meadows with suitable grass species. However, this is rather unlikely to be the case, as *Aptinothrips* are wingless and disperse by wind (Lewis 1973), suggesting that they have little control over the specific meadow which they disperse to. We also found that the performance of asexual *A. rufus* females on *Festuca rubra* positively correlates with the abundance of this grass species at the location the females were collected (Figure 3). This, added to geographical niche variation, suggests that different grass communities favor different clonal assemblages, as different clones perform best on different grass species (van der Kooi et al. 2019, Figure 2). Asexual *Aptinothrips* populations are also genetically highly diverse (van der Kooi and Schwander 2014; Fontcuberta García-Cuenca et al. 2016), meaning that there is a large standing pool of ecologically divergent clones available for selection to act on. Selection would thus result in locally adapted (to the common grasses of their natural environment) asexual thrips populations.

Local adaptation in an asexual but not a closely related sexual species may seem surprising given that asexual species are expected to show less adaptability than their sexual counterparts (Bell 1982). However, depending on the genetic architecture of adaptations to different host plant species, several factors can constrain local adaptation in sexual species. For example, thrips are likely to move (actively or passively) between different grass species within a meadow. Furthermore, *Aptinothrips* are suspected to disperse long distances by wind and human agricultural activities (Lewis 1973). Substantial gene flow between distinct sexual populations might thus impede local adaptation by breaking locally co-adapted gene complexes for host plant selection and use and by introducing mal-adapted alleles into the population (Lenormand 2002). Asexuals could more easily maintain co-adapted gene complexes, and would not be affected by mating with individuals carrying locally mal-adapted gene combinations (Haag and Ebert 2004).

*Aptinothrips* are considered ecological specialists as they exclusively feed on grasses (Palmer 1975). However, within the Poaceae family, thrips display a high degree of generalism, as indicated by the high value of *H’* indices obtained in this study (Figure 2). *A. rufus* displays a small but significant decrease in the level of specialization as compared to *A. elegans*, indicating that it is characterized by a broader ecological niche than the sexual species (Figure 2). Similar to our results in grass thrips, two comparative studies of asexual arthropod species also found that asexuals generally have larger geographic ranges and broader ecological niches than their sexual counterparts (Ross et al. 2013; van der Kooi et al. 2017). At least two (mutually non-exclusive) mechanisms can generate these differences. First, the frozen niche variation model (Vrijenhoek 1979) predicts that when a new clone derives from a sexual ancestor, its phenotypic distribution (and therefore ecological niche) would be narrower than that of its genetically variable sexual ancestor, because a single sexual genotype will be “frozen” in the new asexual clone. However, when multiple clones derive independently from ecologically diverged sexual ancestors, the combination of clones may result in a broad niche at the species level (Vrijenhoek and Parker 2009). Second, broader ecological niches in asexual than sexual species are also predicted by the “General-Purpose Genotype” hypothesis (GPG; Lynch 1984; but see also White 1973; Parker et al. 1977). This hypothesis proposes that asexual clones should have broader niches than sexual individuals because of strong selection for phenotypic plasticity in asexuals. Indeed, a temporally and spatially variable environment should favor, among all the independently derived asexual clones, those with broad environmental tolerances. Our results in asexual *A. rufus* provide support for both mechanisms. We find that there are indeed multiple “more specialist” clones, each with a distinct niche, as predicted by the frozen niche variation model. At the same time, there are also some individual clones with very broad niches, as predicted by the general-purpose genotype hypothesis.

We conclude that niche differentiation between sexual and asexual grass thrips species could locally contribute to their maintenance in sympatry, but not throughout their overlapping distribution ranges. The reproductive advantage of asexual *A. rufus* grass thrips in other locations is striking. Furthermore, it appears that asexuality and a high diversity of ecologically different clones confer *A. rufus* the capacity to colonize and exploit diverse grass communities, which in turn would contribute to maintaining a high genotype diversity in asexual populations. This indicates that sexual reproduction should offer benefits beyond adaptation to different host plants in the thrips, or *A. elegans* will eventually disappear from the grass thrips communities.

## Supporting information

Supplemental: Methods, Figure S1, Tables S1-S3 and References.

## Notes

### Competing Interest Statement

The authors have declared no competing interest.

### Summary of Updates

List of Authors in the manuscript, Table format in manuscript, Abstract first sentence, References formatting, Supplementary Tables Format

