## Supplemental: Methods, Figure S1, Tables S1-S3 and References. for "Ecological specialization and local adaptation in sympatric sexual and asexual grass thrips species"

*Molecular methods for grass community characterization at UNIL, La Sarraz and Blonay*

We verified that the three locations UNIL (geographic coordinates: N 46.52207, E 6.57757), La Sarraz (N 46.65783, E 6.53284), and Blonay (N 46.46759, E 6.91833) were characterized by different grass communities by sampling all grasses in three random plots of 1 $m^2$ at each location. Each plot was divided into a grid of 100 squares (10 x 10 cm each), and the number of squares covered by a given grass morphotype was recorded. The obtained coverages for grass morphotypes were subsequently converted to coverages for grass species using a molecular approach for species determination as follows.

Grass tissue was extracted using the Qiagen plant DNA extraction kit (QIAGEN inc.), following the manufacturer's protocol. We amplified a portion of the mitochondrial maturase K gene, using previously published barcoding primers (matk-390F and matk-1326R; Cuenoud et al. 2002). DNA amplification was done following the Canadian center for DNA barcoding protocol (Fazekas et al. 2012): each reaction contained 2 µl extracted DNA template, 0.75 µl 10× buffer for Platinium Taq (Invitrogen), 1.875 µl 20% trehalose, 0.15 µl dNTP (10 µM), 0.225 µl MgCl2 (50 mM), 0.375 µl of each primer (10 µM), 0.15 µl Taq polymerase (0.75 units, Platinium Taq (Invitrogen)) and were completed to 8.5 µl using nanopure water.

The following cycling conditions were used: 1× 1 min 94° C; 35× 30 sec 94°C, 40 sec 50°C, 40 sec 72°C; 1× 5 min 72°C. 3.5 µl of the PCR-product were run on an ethidiumbromide-stained 1,5% agarose gel. PCR products were purified using ExoSap-

IT according to the manufacturer’s protocol (Isogen Life Science B.V., De Meem, The Netherlands). 5 µl of the purified PCR product was combined with 5 µl forward primer (5 µM) and sent to GATC Biotech, Germany (www.gatc-biotech.com) for sequencing. Obtained sequences were assigned to grass species using BLASTn (blast.ncbi.nlm.nih.gov) with the basic query options. Significant hits with the maximum percentage of identity were recorded as the species of grass present in the field. If the maximum percentage of identity included multiple species, then the species level identification was refined using their distribution range (Lauber et al. 2012). Significant differences in grass composition between locations was tested using a likelihood ratio test for contingency table (Fellows 2012). Log likelihood ratio statistic obtained are the following: (G) = 312.86, X-squared df = 36, p-value $< 2.2e-16$.

*R scripts for LMMs analysis*

```
#setwd("your_directory")
input_file <-"Ecological_specialisation.csv"
data<-read.csv(input_file,  dec=",", sep=";", header=TRUE, na.strings=c("NA"))
attach(data)
nrep=100000
library(lme4)
#LMM global
anovsFtot<-function(rpse,v1,v2,v3,randf,nrep,optimizer){
  obs.F=data.frame(t(anova(lmer(rpse~v1*v2*v3 +
(1|randf),control=lmerControl(optimizer=optimizer)))$F))
  names(obs.F)<-c("v1","v2","v3","v1:v2","v1:v3","v2:v3","v1:v2:v3")
  for (i in 2:nrep) {
```

```
    rpse[randf==1]<-sample(rpse[randf==1])
    rpse[randf==2]<-sample(rpse[randf==2])
    rpse[randf==3]<-sample(rpse[randf==3])
    obs.F[i,]<-t(anova(lmer(rpse~v1*v2*v3                                    +
(1|randf),control=lmerControl(optimizer=optimizer)))$F) }
  pval<-NULL
  for (j in 1:ncol(obs.F)) {
    pval[j]<-sum(obs.F[,j]>=obs.F[1,j])/length(obs.F[,j]) }
  names(pval)<-c("sp","plant","site","sp:plant","sp:site","plant:site","sp:plant:site")
  return(pval)}
pvaltot<-
anovsFtot(data$fit,data$sp,data$plant,data$site,data$batch,nrep,"Nelder_Mead")

#LMM by site

unil<-subset(data,site=="UNIL")
sarraz<-subset(data,site=="La Sarraz")
blonay<-subset(data,site=="Blonay")

anovsFsite<-function(rpse,v1,v2,randf,nrep,optimizer) {
  obs.F=data.frame(t(anova(lmer(rpse~v1*v2+
(1|randf),control=lmerControl(optimizer=optimizer)))$F))
  names(obs.F)<-c("v1","v2","v1:v2")
  for (i in 2:nrep) {
    rpse[randf==1]<-sample(rpse[randf==1])
```

```
    rpse[randf==2]<-sample(rpse[randf==2])
    rpse[randf==3]<-sample(rpse[randf==3])
    obs.F[i,]<-t(anova(lmer(rpse~v1*v2                                                     +
(1|randf),control=lmerControl(optimizer=optimizer)))$F) }
  pval<-NULL
  for (j in 1:ncol(obs.F)) {
    pval[j]<-sum(obs.F[,j]>=obs.F[1,j])/length(obs.F[,j]) }
  names(pval)<-c("sp","plant","sp:plant")
  return(pval)}
pvalunil<-anovsFsite(unil$fit,unil$sp,unil$plant,unil$batch ,nrep,"Nelder_Mead")
pvalsarraz<-anovsFsite(sarraz$fit,     sarraz$sp,     sarraz$plant,     sarraz$batch
,nrep,"Nelder_Mead")
pvalblonay<-anovsFsite(blonay$fit,     blonay$sp,     blonay$plant,     blonay$batch
,nrep,"Nelder_Mead")

#LMM by species
e<-subset(data, sp=="A.elegans")
r<-subset(data, sp=="A.rufus")
anovsFsp<-function(rpse,v1,v2,randf,nrep,optimizer) {
  obs.F=data.frame(t(anova(lmer(rpse~v1*v2+
(1|randf),control=lmerControl(optimizer=optimizer)))$F))
  names(obs.F)<-c("v1","v2","v1:v2")
  for (i in 2:nrep) {
    rpse[randf==1]<-sample(rpse[randf==1])
    rpse[randf==2]<-sample(rpse[randf==2])
```

```
    rpse[randf==3]<-sample(rpse[randf==3])
    obs.F[i,]<-t(anova(lmer(rpse~v1*v2                                    +
(1|randf),control=lmerControl(optimizer=optimizer)))$F) }
  pval<-NULL
  for (j in 1:ncol(obs.F)) {
    pval[j]<-sum(obs.F[,j]>=obs.F[1,j])/length(obs.F[,j]) }
  names(pval)<-c("plant","site","plant:site")
  return(pval)}
pvale<-anovsFsp(e$fit,e$plant,e$site,e$batch,nrep,"Nelder_Mead")
pvalr<-anovsFsp(r$fit,r$plant,r$site,r$batch,nrep,"Nelder_Mead")
#Data for local adaptation of Asexuals A. rufus
input_file2<-"Local_adaptation.csv"
data2<-read.csv(input_file2, header=TRUE, dec=",",sep=";", na.strings=c("NA"))
attach(data2)
poa<-subset(data2,Plant=="P.trivialis")
rub<-subset(data2,Plant=="F.rubra")

#LMM for local adaptation
anovsFadapt<-function(rpse, v1,randf,nrep,optimizer) {
  obs.F=data.frame(t(anova(lmer(rpse~v1+
(1|randf),control=lmerControl(optimizer=optimizer)))$F))
  names(obs.F)<-c("v1")
  for (i in 2:nrep) {
    rpse[randf==1]<-sample(rpse[randf==1])
    rpse[randf==2]<-sample(rpse[randf==2])
```

```
    rpse[randf==3]<-sample(rpse[randf==3])
    rpse[randf==4]<-sample(rpse[randf==4])
    rpse[randf==5]<-sample(rpse[randf==5])
    rpse[randf==6]<-sample(rpse[randf==6])
    rpse[randf==7]<-sample(rpse[randf==7])
    rpse[randf==8]<-sample(rpse[randf==8])
    rpse[randf==9]<-sample(rpse[randf==9])
    rpse[randf==10]<-sample(rpse[randf==10])
    rpse[randf==11]<-sample(rpse[randf==11])
    rpse[randf==12]<-sample(rpse[randf==12])
    rpse[randf==13]<-sample(rpse[randf==13])
    rpse[randf==14]<-sample(rpse[randf==14])
    rpse[randf==15]<-sample(rpse[randf==15])
    obs.F[i,]<-t(anova(lmer(rpse~v1 +
(1|randf),control=lmerControl(optimizer=optimizer)))$F) }
  pval<-NULL
  for (j in 1:ncol(obs.F)) {
    pval[j]<-sum(obs.F[,j]>=obs.F[1,j])/length(obs.F[,j]) }
  names(obs.F)<-c("Coverage")
  return(pval)}
pvalpoa<-
anovsFadapt(poa$Offspring,poa$Poa_trivialis,poa$Date,nrep,"Nelder_Mead")
pvalrub<-
anovsFadapt(rub$Offspring,rub$Festuca_rubra,rub$Date,nrep,"Nelder_Mead")
```

### Supplemental Tables

**Table S1:** Sampling location (ID), altitude (meters), geographic coordinates (CX, CY), locality, number of asexual *A. rufus* females (Nb. replicates) whose performance was measured on *Festuca rubra* and *Poa trivialis*, percentage of the coverage (relative to other grass species) of *Festuca rubra* and *Poa trivialis*. Thrips were sampled from each of these meadows between August and October 2018, and July and October 2019.

| Location | | | | | Nb. replicates | | Grass species coverage (%) | |
|---|---|---|---|---|---|---|---|---|
| ID | Altitude (m) | CX | CY | Locality | *F. rubra* | *P. trivialis* | *F. rubra* | *P. trivialis* |
| 47 | 1362 | N 46.41998989 | E 7.159011333 | Châteaux-d'Oex | 28 | 22 | 36.09 | 0.19 |
| 139 | 765 | N 46.43965323 | E 6.934318607 | Montreux | 7 | 7 | 0.31 | 0 |
| 212 | 841 | N 46.43022091 | E 6.93224917 | Montreux | 30 | 30 | 16.06 | 16.06 |
| 223 | 1004 | N 46.35136259 | E 7.03837334 | Ormont-Dessous | - | 9 | 4.3 | 4.3 |
| 243 | 1516 | N 46.50818257 | E 7.180692113 | Rougemont | - | 7 | 15.29 | 38.23 |
| 254 | 500 | N 46.22537463 | E 7.011018934 | Bex | 33 | 33 | 0 | 35.2 |
| 258 | 540 | N 46.23030987 | E 7.013695817 | Bex | 33 | 33 | 3.06 | 15.32 |
| 284 | 528 | N 46.2378517 | E 7.021140656 | Bex | 33 | 34 | 0.29 | 21.55 |
| 288 | 479 | N 46.46385873 | E 6.866818625 | St-Légier-La Chiésaz | 31 | 32 | 0 | 24.92 |
| 305 | 702 | N 46.45334066 | E 6.910250097 | Montreux | 16 | 22 | 0 | 30.86 |
| 313 | 917 | N 46.44849308 | E 6.939537278 | Montreux | 22 | 15 | 22.46 | 0.3 |
| 618 | 832 | N 46.50448696 | E 6.90040242 | St-Légier-La Chiésaz | 30 | 33 | 21.61 | 0.29 |
| 626 | 841 | N 46.46731851 | E 7.047137253 | Rossinière | 16 | - | 48.39 | 0 |
| 630 | 985 | N 46.48100467 | E 7.179806989 | Châteaux-d'Oex | 7 | - | 61.98 | 0 |
| 636 | 964 | N 46.45815679 | E 6.923316457 | Montreux | 34 | 16 | 60.18 | 0.02 |
| 891 | 1400 | N 46.51026655 | E 7.195655416 | Rougemont | 35 | 30 | 90.8 | 0 |

**Table S2:** Details of the LMMs used to assess the variation of performances (number of offspring produced) according to thrips species (*A. elegans* or *A. rufus*) the grass species (seven species tested, see main text) and the three thrips sampling locations. F-value is the observed value, p-values are estimated based on 100’000 randomizations as described in the methods of the main text. Significant p-values are indicated in bold.

| | | Data total | | Data split by Location | | | | | | Data split by species | | | |
|---|---|---|---|---|---|---|---|---|---|---|---|---|---|
| | | | | UNIL | | La Sarraz | | Blonay | | *A. elegans* | | *A. rufus* | |
| Explanatory variable | df | F-value | P-value | F-Value | P-value | F-value | P-value | F-value | P-value | F-Value | P-value | F-value | P-value |
| Thrips sp. | 1 | 99.2 | **0.00001** | 9.9 | **0.0037** | 23.7 | **0.00001** | 84.6 | **0.00001** | - | - | - | - |
| Grass sp. | 6 | 8.1 | **0.00001** | 3.3 | **0.00478** | 5.4 | **0.00014** | 2.4 | **0.03523** | 13 | **0.00001** | 3.1 | **0.00652** |
| Location | 2 | 3.6 | **0.02668** | - | - | | - | - | - | 3.8 | **0.01624** | 9.5 | **0.00012** |
| Thrips sp. : Grass sp. | 6 | 2.4 | **0.02993** | 2.8 | **0.01242** | 2.3 | **0.04054** | 1.8 | 0.11086 | - | - | - | - |
| Thrips sp. : Location | 2 | 12.4 | **0.00001** | - | - | - | - | - | - | - | - | - | - |
| Grass sp. : Location | 12 | 1.2 | 0.25781 | - | - | - | - | - | - | 1.2 | 0.26354 | 1.8 | **0.04973** |
| Thrips sp. : Grass sp: Location | 12 | 2.1 | **0.01616** | - | - | - | - | - | - | - | - | - | - |

**Table S3:** Details of the LMMs used to assess the association between the performance of *A. rufus* females and host plant coverage across 16 meadows. F-value is the observed value, p-values are estimated based on 100’000 randomizations across meadows as described in the methods of the main text. Significant p-values are indicated in bold.

| | | Grass species | | | |
|---|---|---|---|---|---|
| | | F. rubra | | P. trivialis | |
| Explenatory variable | df | F-value | P-value | F-value | P-value |
| Coverage | 13 | 26.23302 | **0.00187** | 0.0154398 | 0.94346 |

**Supplemental Figures**

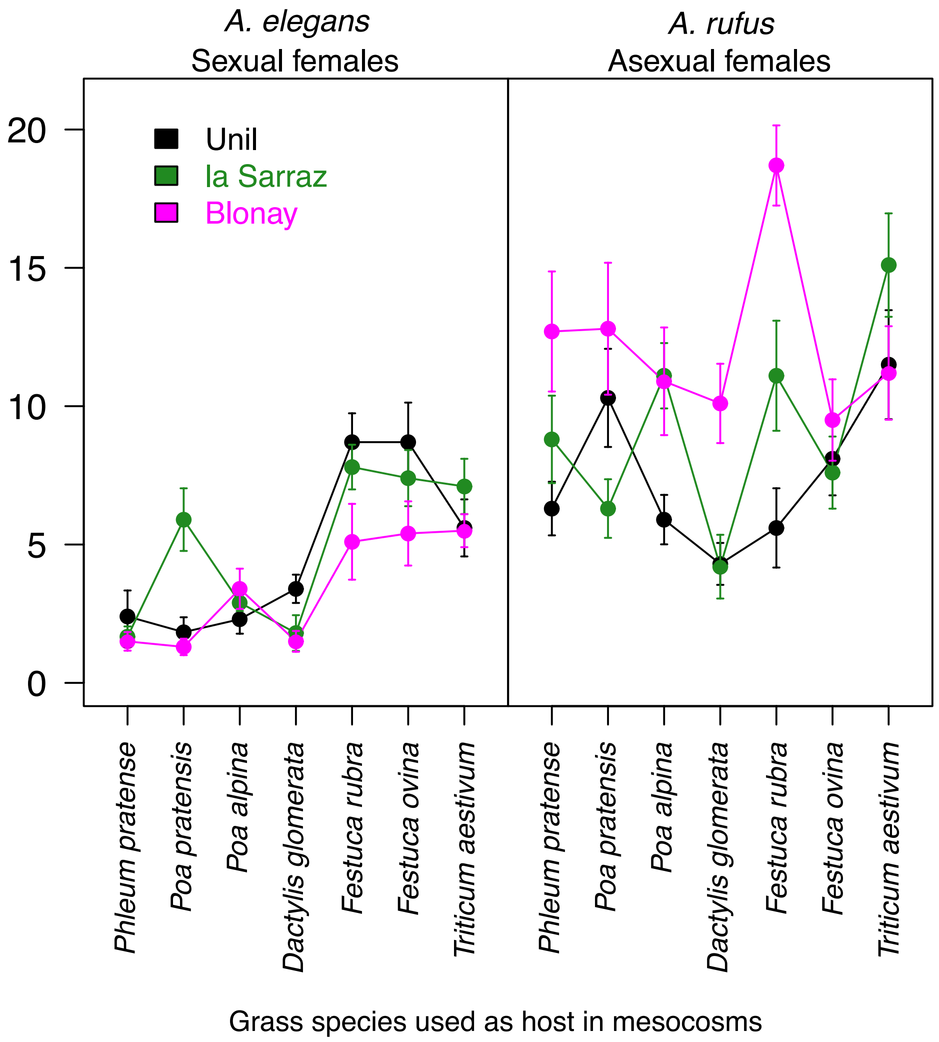


**Figure S1:** Average performance on different grass hosts (number of offspring produced) and standard deviation for the two thrips species collected at different locations. The data used are the same as in Figure 1 in the main text but separated by species to illustrate performance differences and similarities between thrips from different locations. The lines connecting points have no biological meaning but are added for a better visualization.
